# Zika virus alters centrosome organization to suppress the innate immune response

**DOI:** 10.1101/2020.09.15.298083

**Authors:** Andrew Kodani, Kristeene A. Knopp, Elizabeth Di Lullo, Hanna Retallack, Arnold R. Kriegstein, Joseph L. DeRisi, Jeremy F. Reiter

## Abstract

Zika virus (ZIKV) is a flavivirus transmitted via mosquitoes and sex to cause congenital neurodevelopmental defects, including microcephaly. Inherited forms of microcephaly (MCPH) are associated with disrupted centrosome organization. Similarly, we found that ZIKV infection disrupted centrosome organization. ZIKV infection disrupted the organization of centrosomal proteins including CEP63, a MCPH-associated protein. The ZIKV nonstructural protein NS3 bound CEP63, and expression of NS3 was sufficient to alter centrosome architecture and CEP63 localization. Loss of CEP63 suppressed ZIKV-induced centrosome disorganization, indicating that ZIKV requires CEP63 to disrupt centrosome organization. ZIKV infection or loss of CEP63 decreased the centrosomal localization and stability of TANK-binding kinase 1 (TBK1), a regulator of the innate immune response. ZIKV infection or loss of CEP63 also increased the centrosomal accumulation of the CEP63 interactors, Mindbomb1 (MIB1) and DTX4, ubiquitin ligases that respectively activate and degrade TBK1. Therefore, we propose that ZIKV NS3 binds CEP63 to increase centrosomal DTX4 localization and destabilization of TBK1, thereby tempering the innate immune response. In addition to identifying a mechanism by which CEP63 controls the innate immune responses in ZIKV infection, we propose that the altered centrosomal organization caused by altered CEP63 function may contribute to ZIKV-associated microcephaly.

## Introduction

In the summer of 2015, cerebral malformations were linked to mosquito-transmitted Zika virus (ZIKV). Inherited forms of microcephaly (MCPH) are characterized by reduced head and brain size, resulting in severe intellectual disability and motor movement defects. Many forms of MCPH are caused by autosomally recessive mutations in genes encoding centrosomal proteins required for centrosome biogenesis and mitotic progression ^1^. Given the similar pathologies between ZIKV-associated and inherited MCPH, we hypothesized that both disorders were due to centrosomal defects leading to disrupted brain development.

In mammalian cells, centrosomes serve as the microtubule organizing center of the cell to facilitate neuronal migration and cell division ^2-4^. The centrosome is composed of centrioles surrounded by a pericentrosomal matrix that nucleates microtubules ^5,6^. During S phase, the centrosome duplicates by recruiting specialized proteins to the centriole base ^5,7-9^. Many MCPH-associated proteins are recruited to the centrosome in a hierarchical manner to promote centrosome duplication ^7,10-13^. Defects in centrosome organization and biogenesis leading to cell death or premature differentiation in neural progenitors may underlie the pathology of many forms of MCPH ^10,14,15^.

Cells infected with ZIKV have disrupted centrosome organization and mitotic abnormalities, leading to altered neural progenitor differentiation ^16-20^. However, the mechanism by which ZIKV disrupts centrosome architecture remains unclear. We found that ZIKV alters the function of the MCPH-associated protein, CEP63. More specifically, the ZIKV nonstructural protein, NS3, localizes to the centrosome and binds CEP63. ZIKV NS3 overaccumulates CEP63 at centrosomes of ZIKV-infected cells and recruits the ubiquitin ligases, MIB1 and DTX4. MIB1 and DTX4 promote the degradation of TBK1, a regulator of the innate immune response ^21-25^. Consequently, ZIKV-infected cells express lower levels of interferon β (IFNβ), a key anti-viral signal. We propose that ZIKV recruits CEP63 to the centrosome to degrade TBK1, dampening anti-viral responses, and disrupted centrosomal architecture in ZIKV-infected cells may also perturb neural development.

## Results

### ZIKV infection disorganizes the centrosome

To explore whether microcephaly associated with ZIKV infection involves the centrosome, we infected human induced neural precursor cells (iNPCs) derived from induced pluripotent stem cells with ZIKV and examined the organization of their centrosomes. Sixteen hours post infection (hpi) with ZIKV, iNPCs displayed supernumerary foci of Centrin, a component of the centriolar distal lumen ^26^ (**Figure 1A, C**). Similarly, ZIKV-infected U87 and H4 cells exhibited supernumerary Centrin foci (**Figure 1B, C,** and Supplemental Figure 1A). Like ZIKV-infected cultured cells, NPCs isolated from human fetal neocortical tissue infected by ZIKV exhibited supernumerary Centrin foci (**Figure 1D**).

**Figure 1:**
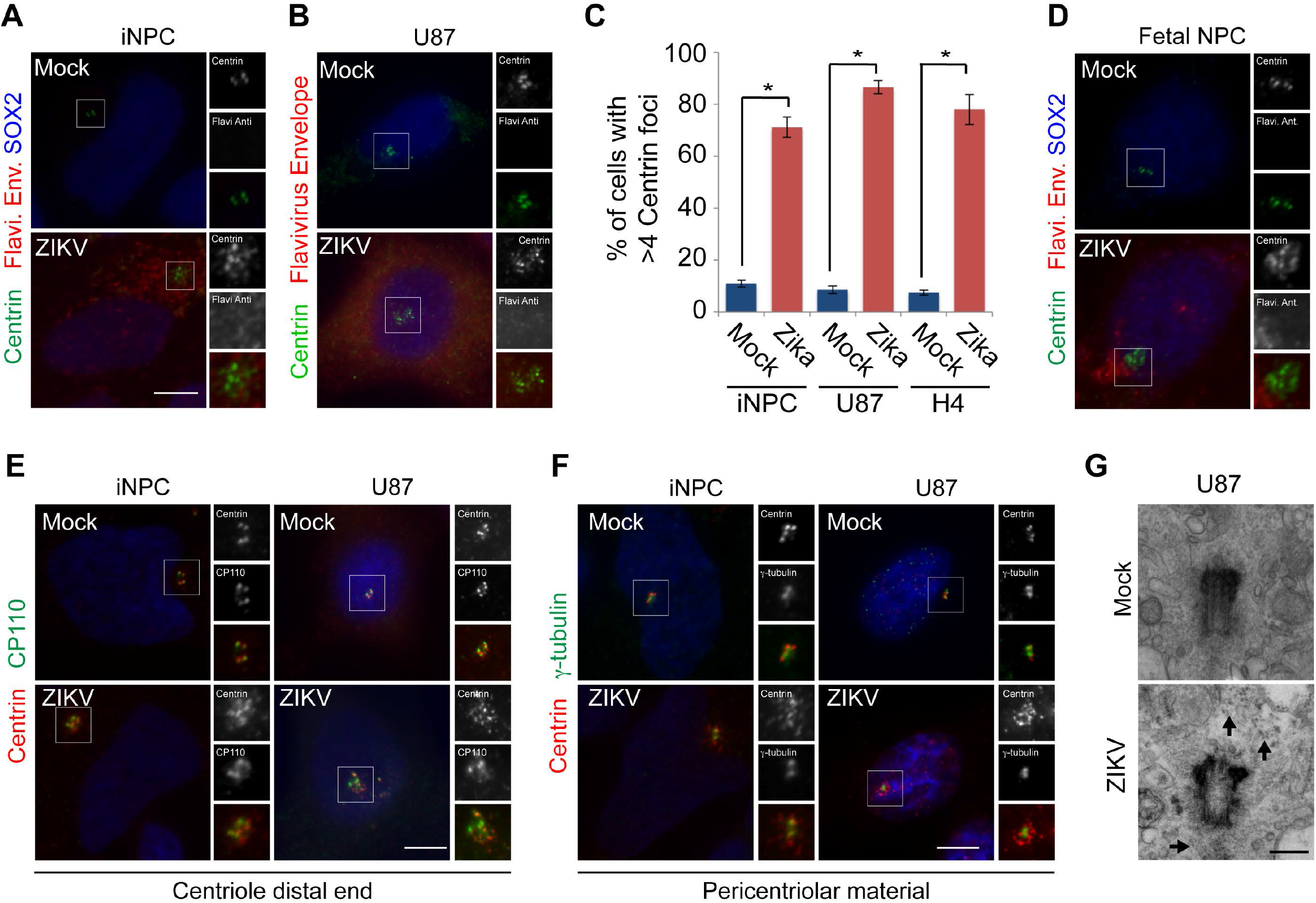
ZIKV infection induces the formation of supernumerary Centrin foci in neural precursor cells. **(A)** Neural precursors derived from human iPS cells (iNPCs) were mock or ZIKV-infected and co-stained for Centrin (green), Sox2 (blue) and flavivirus envelope (red). All cells were fixed 16 hours post infection (hpi). **(B)** Mock and ZIKV-infected U87 and H4 cells in S phase co-stained for Centrin (green), flavivirus envelope (red) and DNA (blue). **(C)** Quantification of the percentage of mock or ZIKV-infected iNPC, U87 or H4 cells in S phase with greater than four Centrin foci. Asterisk denotes p<0.005 (Student’s t test) **(D)** Neural precursor cells mock or ZIKV-infected and co-stained for Centrin (green), flavivirus envelope (red) and SOX2 (blue). **(E)** Mock or ZIKV-infected iNPC and U87 cells in S phase co-stained for Centrin (red) and the centriole distal end component, CP110 (green). **(F)** Mock or ZIKV-infected iNPC and U87 cells co-stained for Centrin (red) and the pericentriolar material proteins γ-tubulin (green) **(G)** Transmission electron micrographs of centrioles from mock or ZIKV-infected U87 cells. Scale bars indicate 100nm. Arrows indicate electron dense particles.

In ZIKV-infected iNPCs and U87 cells, the supernumerary Centrin foci co-localized with the distal centriole protein CP110 ^27^ (**Figure 1E** and Supplemental Figure 1B), indicating that ZIKV-associated supernumerary Centrin foci can recruit centriolar proteins. However, the supernumerary Centrin foci do not accumulate the distal appendage protein CEP164 ^28^ or pericentrosomal protein γ-tubulin ^29^ (**Figure 1F,** Supplementary Figure 1B, C). ZIKV infection did not alter the levels of CP110 or γ-tubulin (Supplemental Figure 1D). Electron microscopy revealed that ZIKV infection did not alter centriole number or distal appendages 16 hpi (**Figure 1G**). Instead, ZIKV-infected cells accumulated electron dense particles in the vicinity of the centrosome. Thus, acute ZIKV infection does not cause centrosome overproduction, but rather disorganizes the centrosome.

### ZIKV disrupts centrosome organization in a CEP63-dependent manner

Many MCPH-associated proteins dynamically localize to the centrosome during its biogenesis ^30-32^. Given that ZIKV infection alters centrosome organization, we investigated whether the localization of MCPH-associated proteins is also altered. ZIKV infection in U87 cells did not affect either the localization or levels of the MCPH-associated proteins PLK4, STIL, SAS6, CDK5RAP2, CEP152, or WDR62 (Supplemental Figure 2A-D). In striking contrast, ZIKV infection caused CEP63, another MCPH-associated protein ^7^, to overaccumulate at supernumerary Centrin foci (**Figure 2A**, and quantified in **Figure 2B**). As in U87 cells, ZIKV infection of iNPCs and NPCs isolated from dissociated fetal brain tissue relocalized CEP63 to supernumerary Centrin foci (**Figure 2C**). Although ZIKV infection did not dramatically affect CEP63 protein levels, it did increase the levels of a higher molecular weight species of CEP63 (**Figure 2D**).

**Figure 2:**
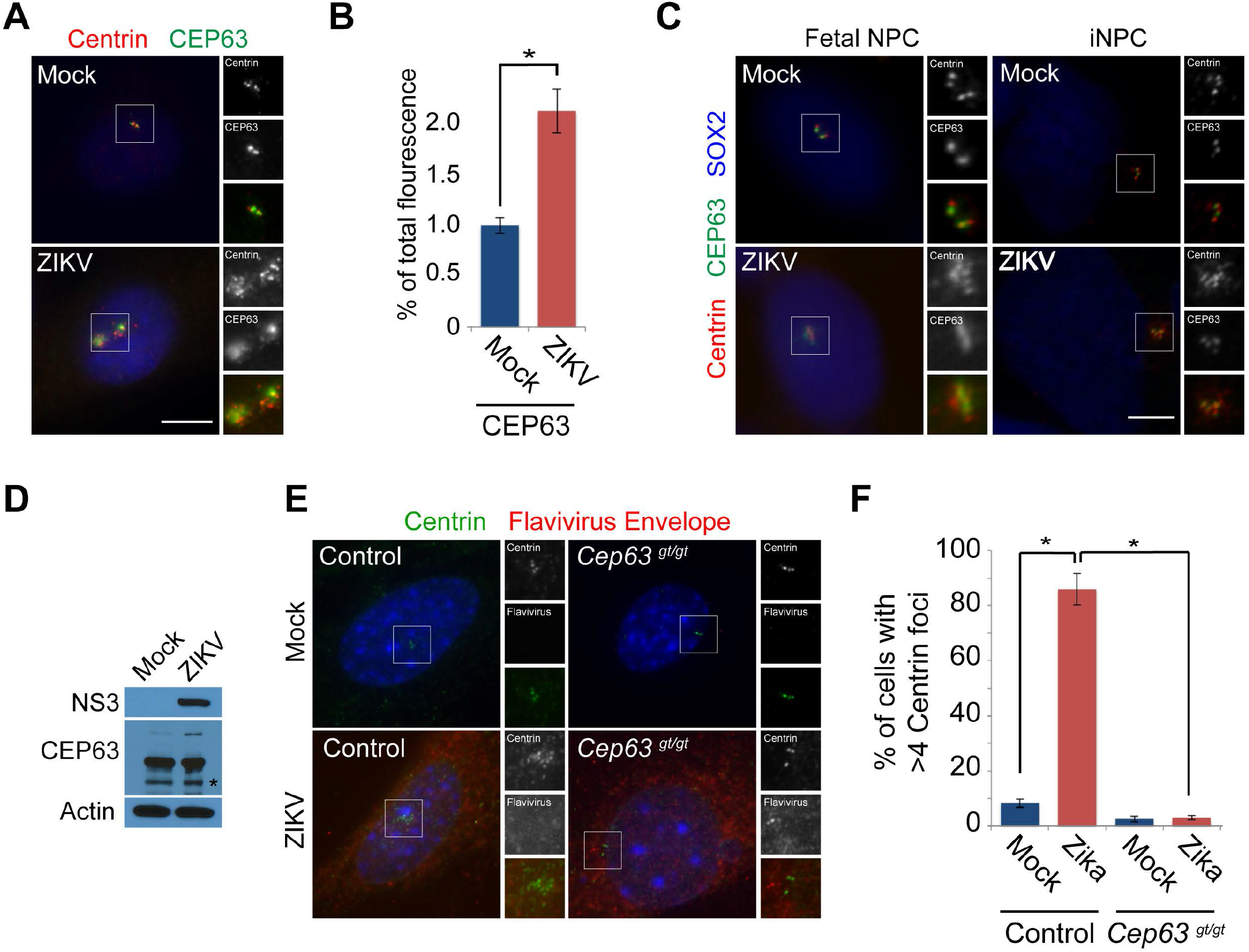
ZIKV induces supernumary Centrin foci in a CEP63-dependent manner. **(A)** Mock or ZIKV-infected U87 cells in S phase co-stained for Centrin (red) and the centrosomal microcephaly-associated protein, CEP63 (green). **(B)** The fluorescence intensity ± s.d. of CEP63 were quantified in mock and ZIKV infected U87 cells. For fluorescence quantifications 8 cells were analyzed per experiment (n=3). Asterisk denotes p<0.005 (Student’s t test). **(C)** Mock and ZIKV-infected neuronal progenitors (NPC) isolated from fetal brain cortex and iPSC derived neural precursor cells (iNPC) co-stained for Centrin (red), CEP63 (green) and SOX2 (blue). **(D)** Immunoblot analysis of lysate from mock or ZIKV-infected U87 cells probed for CEP63 and ZIKV NS3. Actin served as a loading control. Asterisk denotes specific band. **(E)** S-phase mock and ZIKV infected control and *Cep63*^gt/gt^ MEFs co-stained for Centrin (green) and flavivirus envelope (red). Scale bars indicate 5μM for all images. All cells were fixed 16 hours post infection (hpi), **(F)** Quantification of the percentage of mock or ZIKV-infected Het and *Cep63*^gt/gt^ MEFs in S phase with greater than four Centrin foci. Asterisk denotes p<0.005 (Student’s t test).

Because MCPH-associated proteins function in centrosome biogenesis, we hypothesized that the disruption of CEP63 localization participates in ZIKV-associated centrosomal disorganization. To test this, we infected control and *Cep63^gt/gt^* mouse embryonic fibroblasts (MEFs) ^8^ with ZIKV and assessed Centrin organization. In contrast to control cells, ZIKV did not induce supenumerary Centrin foci in the absence of Cep63 (**Figure 2E-F**). Together, these findings indicate that CEP63 is required for ZIKV-associated reorganization of the centrosome.

### ZIKV NS3 binds CEP63 to disrupt centrosome organization

In ZIKV-infected iNPCs, the non-structural viral protein NS3 localized to one or two foci among the supernumerary Centrin foci at 16hpi (**Figure 3A**). A yeast two-hybrid analysis suggested that CEP63 binds the non-structural protein NS3 of flaviviruses related to ZIKV ^33^. As ZIKV infection causes CEP63 to overaccumulate at centrosomes, we determined whether ZIKV NS3 interacts with CEP63. Immunoprecipitation of endogenous CEP63 in ZIKV-infected cells revealed that CEP63 interacts with ZIKV NS3 (**Figure 3B**). To confirm the interaction of ZIKV NS3 with CEP63, we transfected cells with Myc-tagged Brazilian ZIKV NS3 and found that Myc-NS3 co-precipitated with endogenous CEP63, but not CP110, another centrosomal protein that misaccumulates at the centrosome upon ZIKV infection (**Figure 3C**).

**Figure 3:**
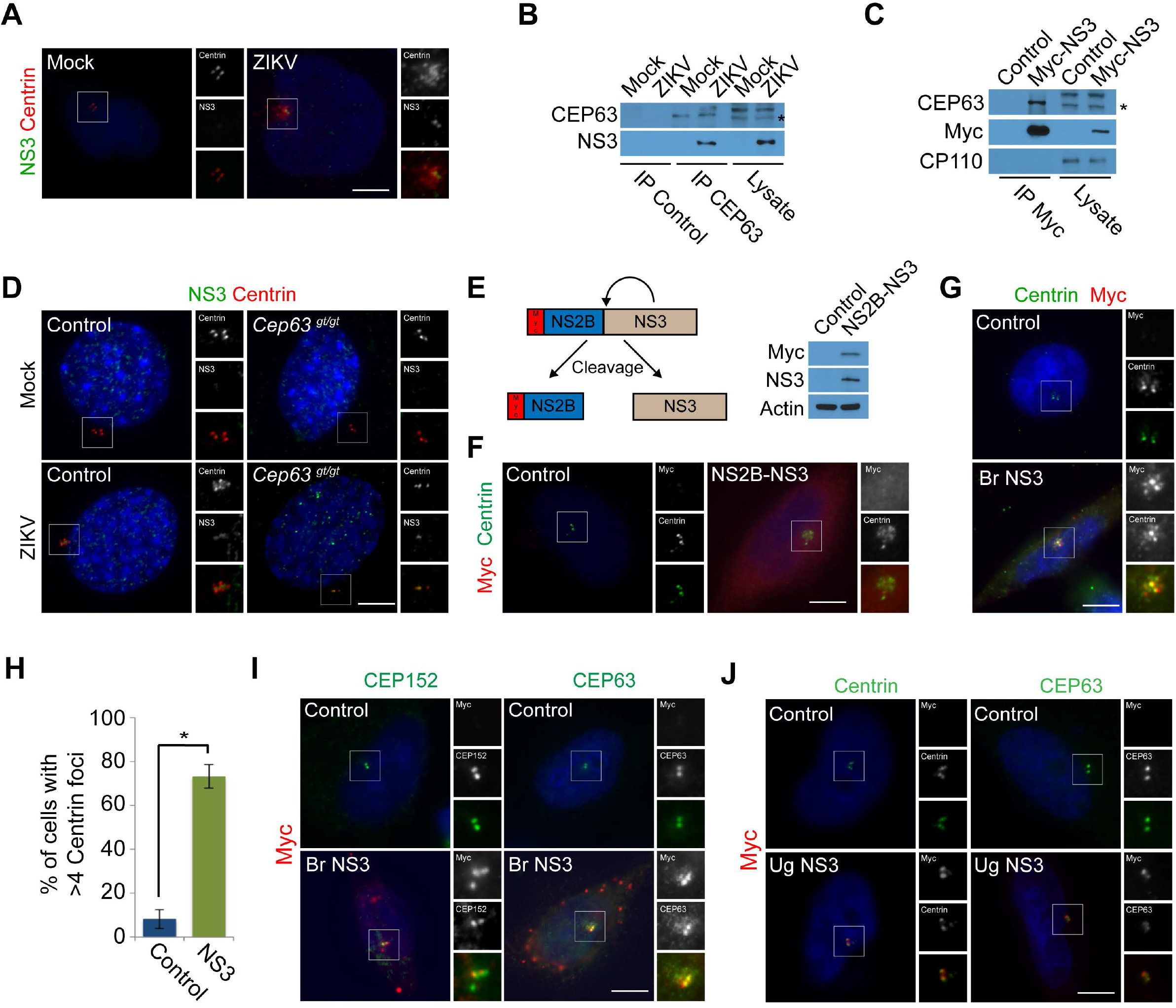
ZIKV NS3 binds the microcephaly-associated protein CEP63 and accumulates it at the centrosome. **(A)** Mock or ZIKV-infected iNPC cells were co-stained for Centrin (red) and NS3 (green). (**B)** Mock or ZIKV-infected U87 cell lysates were immunoprecipitated for CEP63 or c-Myc, which served as a negative control for this and other immunoprecipitations (Control). Precipitated proteins were immunoblotted for CEP63 and NS3. Asterisk denotes specific band. **(C)** We expressed Myc-tagged Brazilian Zika NS3 in 293T/17 cells and immunoprecipitated c-Myc. Precipitated proteins were probed for the Myc tag of NS3 and endogenous CEP63. The centriole distal end component, CP110 served as a negative control. **(D)** Mock and ZIKV-infected control and *Cep63*^gt/gt^ MEFs were co-stained for Centrin (red) and ZIKV NS3 (green). **(E)** Schematic representation of Myc-NS2B-NS3 being proteolytically cleaved by itself to generate Myc-NS2B and NS3 proteins. Immunoblot of HeLa cells transfected with Myc-NS2B-NS3 probed with antibodies to Myc and NS3. Actin served as a loading control. **(F)** Mock or Myc-NS2B-NS3 expressing HeLa cells co-stained for c-Myc (red) and Centrin (green). **(G)** Quantification of the percentage of control or NS2B-NS3 expressing cells in S-phase with greater than four Centrin foci. Asterisk denotes p<0.005 (Student’s t test). **(H)** Quantification of the percentage of control and NS3 expressing HeLa cells with greater than four Centrin foci. For all quantifications at least 100 cells were counted per experiment (n=3). p<0.005 (paired Student’s t test) for the comparison of NS3 to mock transfected cells was denoted with asterisk. **(I)** Control or Myc-tagged Brazilian ZIKV NS3 (Br NS3) transfected HeLa cells were co-stained for Myc (red) and MCPH protein CEP152 or CEP63. **(J)** Control or Myc-tagged Ugandan NS3 (Ug NS3) expressing HeLa cells were co-stained for Myc (red) and Centrin (green) or CEP63 (green). Scale bars indicate 5μM for all images.

As ZIKV NS3 interacts with CEP63, we investigated whether CEP63 was required to localize NS3 to the centrosome. To test this, we infected control and *Cep63^gt/gt^* MEFs with ZIKV and examined NS3 localization 16 hpi. Similar to control cells, ZIKV NS3 localized to centrosomes in *Cep63^gt/gt^* MEFs (**Figure 3D)** suggesting that NS3 affects CEP63 localization, but not vice versa.

NS3 acts together with another nonstructural protein, NS2B, to form a proteolytic complex **(Figure 3E)** ^34^. To gain insight into how ZIKV disrupts centrosomes, we examined whether the NS2B/NS3 complex could induce supernumerary Centrin foci or if NS3 alone is sufficient. Expression of either Brazilian ZIKV Myc-NS2B/NS3 or Myc-NS3 induced supernumerary Centrin foci (**Figure 3F-H**), suggesting NS3 alone is sufficient to perturb centrosome organization. Additionally, Myc-NS3 increased the centrosomal accumulation of CEP63 (**Figure 3I** and quantified in Supplementary Figure 3D) and had no effect on the localization or stability of CEP152, CDK5RAP2 and WDR62 (**Figure 3I** and Supplemental Figure 3B-D). Therefore, Brazilian ZIKV NS3 interacts with CEP63, localizes to the centrosome, and is sufficient to reorganize centrosomal architecture akin to ZIKV infection.

Microcephaly is associated with South American strains of ZIKV, but not Uganda ZIKV ^35-37^ Given the NS3 proteins of the Brazilian and Ugandan strains of ZIKV differ by 11 amino acids, we examined whether Ugandan NS3 was sufficient to induce supernumerary Centrin foci and misrecruit CEP63. Like Brazilian NS3, Ugandan ZIKV NS3 localized to centrosomes (**Figure 3J)**. In striking contrast to Brazilian ZIKV NS3, the Ugandan NS3 did not induce supernumerary Centrin foci or increase centrosomal CEP63 localization (**Figure 3J,** and quantified in Supplementary Figure 3E, F). Together, these findings indicate that Brazilian ZIKV NS3 has acquired the ability to increase centrosomal CEP63 localization and induce supernumerary Centrin foci.

### ZIKV suppresses innate immune signaling in a CEP63-dependent manner

To assess how centrosomal disorganization participates in ZIKV biology, we examined whether centrosomal disorganization contributes to viral production. We found that loss of CEP63 did not alter ZIKV production (**Figure 4A**).

**Figure 4:**
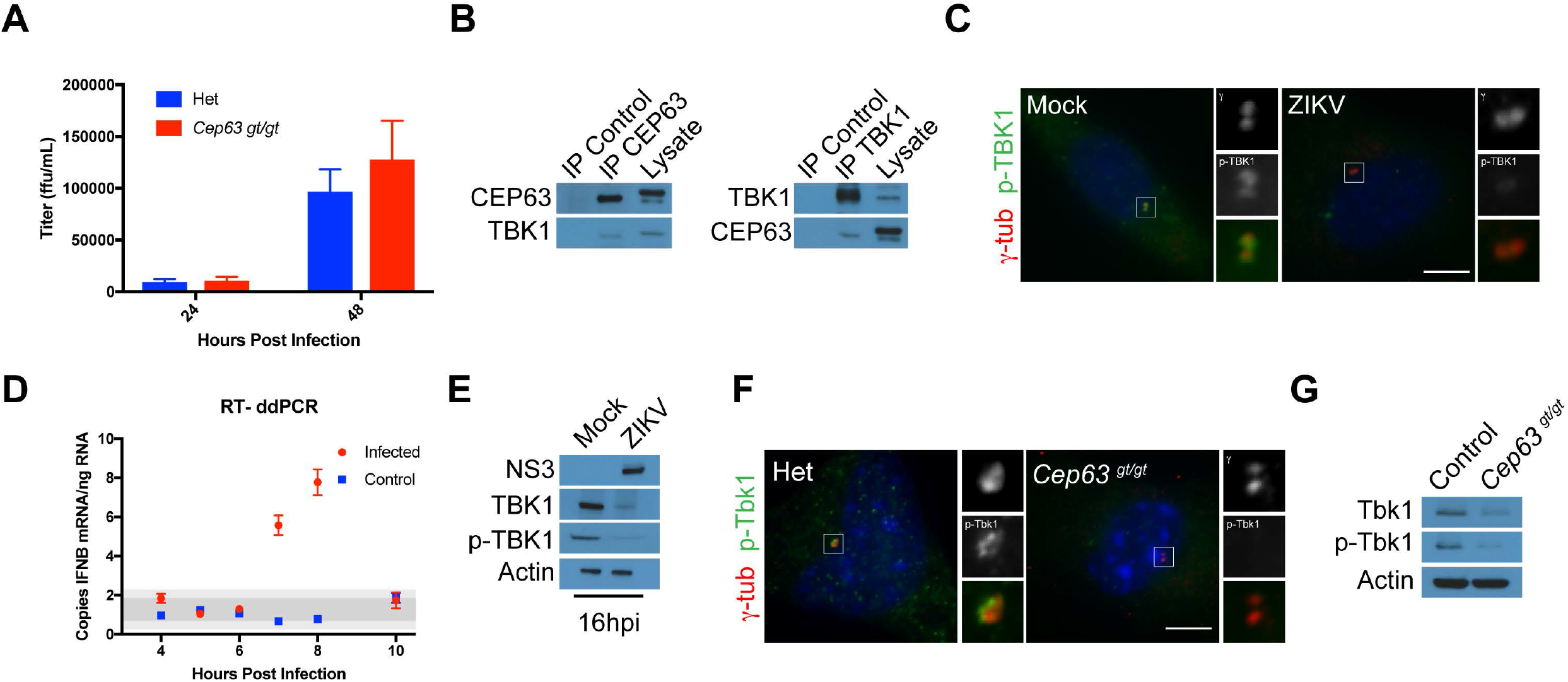
ZIKV disrupts the innate immune response. **(A)** Viral titers from Control and *Cep63^gt/gt^* MEFs at 24, and 48 hpi. There was no statistical difference between control and *Cep63^gt/gt^* viral titers at the two time points (paired student’s t-test). **(B)** HeLa cell lysates were immunoprecipitated with antibodies to CEP63 and TBK1. Precipitated proteins were immunoblotted for CEP63 and TBK1. **(C)** Mock and ZIKV-infected U87 cells were co-stained for γ-tubulin (red) and p-TBK1 (green). **(D)** RT-ddPCR of IFNβ transcript at the indicated time points in mock and ZIKV-infected H4 cells. **(E)** Lysate from mock or ZIKV-infected U87 cells at 16 hpi were immunoblotted with antibodies to NS3, TBK1, and p-TBK1. Actin served as a loading control. **(F)** Control and *Cep63*^gt/gt^ MEFs were co-stained with antibodies to γ-tubulin (red) and p-Tbk1 (green). Scale bars indicate 5μM for all images. **(G)** Immunoblot analysis of control and *Cep63*^gt/gt^ MEFs probed with antibodies to Tbk1 and p-Tbk1. Actin served as a loading control.

As CEP63 is not required for ZIKV production, we hypothesized that it may alter the cellular response to ZIKV infection. ZIKV inhibits type I interferon effector signaling by degrading STAT2, and by an unknown mechanism, inhibits interferon induction ^38,39^. During viral infections, the kinase TBK1 phosphorylates and activates the transcription factor IRF3, and is then degraded, to regulate interferon induction ^23,40,41^. TBK1 localizes to centrosomes ^42-47^ and proximity interaction studies have suggested that TBK1 and CEP63 interact ^46^. Through co-immunoprecipitation, we confirmed that endogenous CEP63 and TBK1 interact (**Figure 4B**). An activated form of TBK1, phospho-TBK1 (p-TBK1), is removed from centrosomes upon ZIKV infection during mitosis ^47^, we confirmed p-TBK1 is removed during interphase (**Figure 4C**, and quantified in Supplemental Figure 4A).

As p-TBK1 is absent from centrosomes of ZIKV-infected cells, we assessed whether ZIKV infection induces interferon beta (IFNβ) using reverse-transcription digital droplet PCR (RT-ddPCR). Following ZIKV infection, IFNβ expression gradually peaked 6 to 8 hpi, (**Figure 4D**). IFNβ expression was abruptly curtailed at 8 to 10 hpi in ZIKV-infected cells, corresponding to the timing of when centrosome begin to disorganize. ZIKV infection decreased levels of TBK1 and phosphorylated TBK1 (p-TBK1) at 16 hpi (**Figure 4E**). Therefore, we propose that ZIKV may suppress the innate immune response by suppressing TBK1 stability and centrosome organization.

To test whether ZIKV disrupts TBK1 function in a CEP63-dependent manner, we examined whether centrosomal p-TBK1 localization depends upon CEP63. Indeed, p-TBK1 was absent from centrosomes in *Cep63^gt/gt^* MEFs (**Figure 4F**, and quantified in Supplemental Figure 4B), indicating that CEP63 is required for the centrosomal localization of p-TBK1. Similarly, TBK1 and p-TBK1 levels were decreased in *Cep63^gt/gt^* MEFs **(Figure 4G)**. These data are consistent with ZIKV interfering with CEP63 function to disrupt TBK1 stability.

### ZIKV infection increases centrosomal MIB1 and DTX4 to degrade TBK1

How might ZIKV reorganize the centrosome in a CEP63-dependent manner to affect TBK1 stability? In response to Sendai virus infections, Mindbomb1 (MIB1) binds and activates TBK1 through K63-linked ubiquitylation ^21,25^, and proximity interaction studies of CEP63 identified MIB1 as an interactor ^46^. We confirmed the interaction of MIB1 to TBK1 and CEP63 using reciprocal co-immunoprecipitation of endogenous MIB1 with TBK1 and CEP63 (**Figure 5A** and Supplemental Figure 5A). Furthermore, we assessed whether ZIKV infection induced a change in the centrosomal localization of MIB1. ZIKV infection increased centrosomal accumulation of MIB1 in iNPCs, fetal NPCs, and U87 cells (**Figure 5B** and Supplemental Figure 5B, C). ZIKV infection mildly altered the stability of MIB1, but significantly increased the levels of a higher molecular weight species (**Figure 5C**).

**Figure 5:**
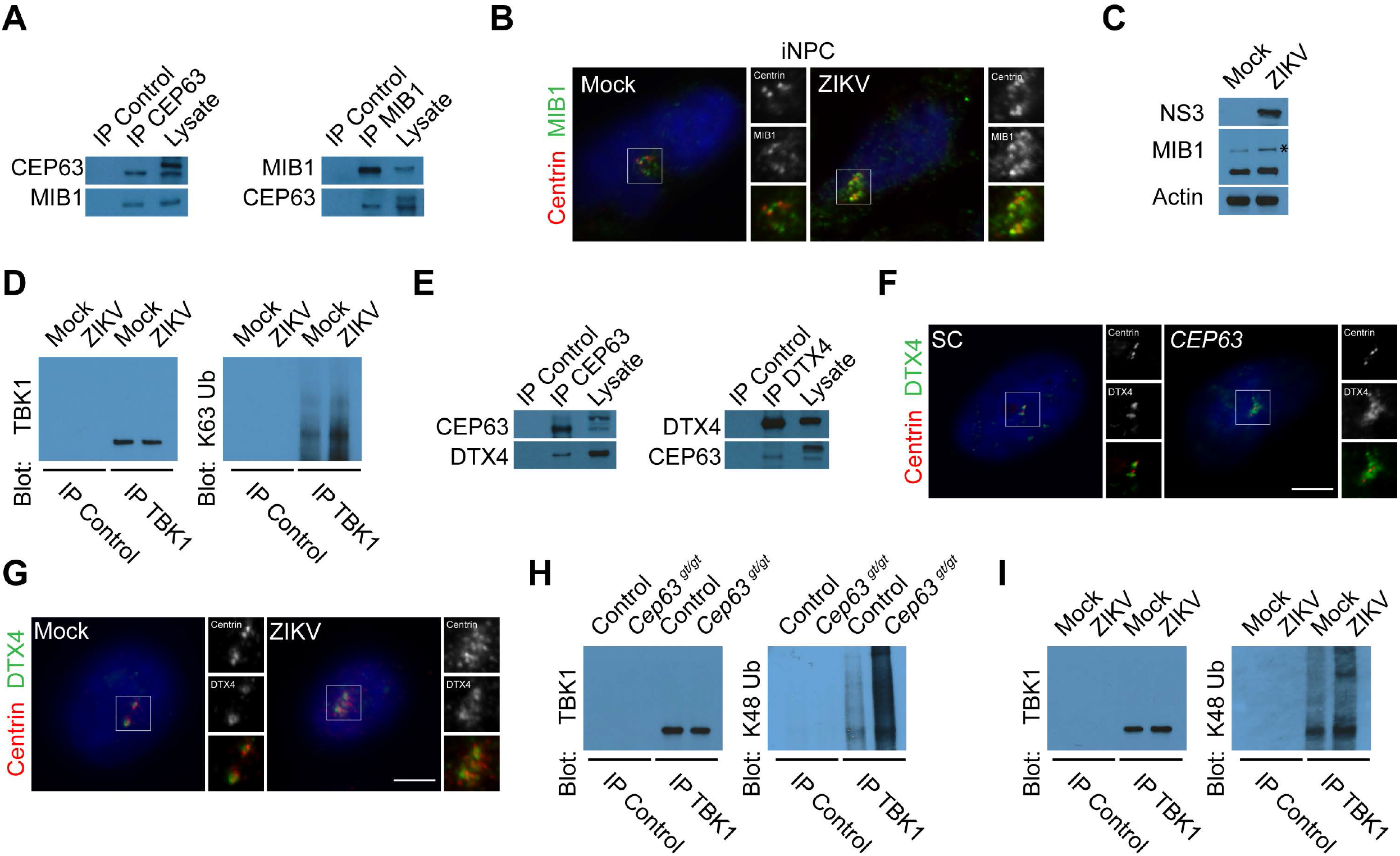
ZIKV increases the ubiquitylation of TBK1 by mislocalizing the ubiquitin ligases MIB1 and DTX4. **(A)** HeLa cell lysate was immunoprecipitated for CEP63, MIB1, or c-Myc (Control). Precipited proteins were immunoblotted for CEP63 and MIB1. **(B)** Mock or ZIKV infected iNPC cells co-stained for Centrin (red) and MIB1 (green). All cells were fixed and stained 16 hpi. **(C)** Immunoblots for mock and ZIKV-infected cell lysates probed for NS3 and MIB1. Actin served as a loading control. Asterisk denotes higher molecular weight species of MIB1. **(D)** Total cell lysates from mock or ZIKV-infected U87 cells 16hpi were immunoprecipitated for TBK1 or c-Myc. The TBK1 immunoprecipitation was probed with TBK1 and K63-linked ubiquitin. **(E)** HeLa cell lysate was immunoprecipitated with antibodies to CEP63, DTX4 or c-Myc (negative control). Precipitated proteins were blotted with antibodies to CEP63 and DTX4. **(F)** SC and *CEP63* siRNA treated HeLa cells were co-stained for Centrin (red) and DTX4 (green). **(G)** Mock and ZIKV-infected U87 cells costained for Centrin (red) and DTX4 (green). Scale bars indicate 5μM for all images. **(H)** TBK1 or c-Myc (negative control) were immunoprecipitated from Control or *Cep63* mutant MEFs. Precipitat proteins were probed with antibodies to TBK1 and K48-linked ubiquitin. **(I)** 16hpi Mock and ZIKV-infected U87 cell lysate was immunoprecipitated for TBK1 or c-Myc. The immunoprecipitations were probed using antibodies to TBK1 and K-48-linked ubiquitin.

Because CEP63 function is altered in ZIKV infected cells and CEP63 interacts with MIB1, we assessed whether CEP63 restricts MIB1 localization to the centrosome. Like ZIKV infection, depletion of CEP63 increased MIB1 accumulation at the centrosome and resulted in the appearance of a higher molecular weight species of MIB1 (Supplemental Figure 5D-F), indicating that MIB1 localization to centrosomes is restricted by CEP63.

We then tested whether CEP63 regulates MIB1 activity by examining the levels of MIB1-dependent K63-ubiquitylation of TBK1 in *Cep63^gt/gt^* MEFs. K63-ubiquitylation of TBK1 was significantly increased in *Cep63* mutant cells, suggesting that increased centrosomal MIB1 may facilitate its ability to ubiquitylate TBK1 at the centrosome (Supplemental Figure 5G-H). Similar to the loss of CEP63, K63-ubiquitylation of TBK1 was dramatically increased in ZIKV infected cells **(Figure 5D** and Supplemental Figure 5I). Together, these data suggest that ZIKV disrupts CEP63 to affect the levels and function of MIB1 at the centrosome.

Once activated the innate immune response is attenuated by K48 ubiquitin-mediated degradation of TBK1 by the ubiquitin ligase DTX4 ^41^. As TBK1 localizes to the centrosome, we examined whether DTX4 is, like MIB1, present at centrosomes. Interestingly, DTX4 partially co-localized with γ-tubulin (Supplemental Figure 5J). The specificity of the antibody was confirmed by siRNA (Supplemental Figure 5J). In accordance with previously published data, depletion of DTX4 led to the stabilization of TBK1 ^41^ (Supplemental Figure 5K).

As TBK1 stability is disrupted in the absence of CEP63, we hypothesized that DTX4 interacts with and is localized to the centrosome by CEP63. Reciprocal immunoprecipitation of CEP63 and DTX4 confirmed that these two proteins interact (**Figure 5E**). Similar to MIB1, DTX4 accumulated in the vicinity of the centrosome in *CEP63*-depleted and ZIKV-infected cells (**Figure 5F-G,** Supplemental Figure 5L-M), suggesting that DTX4 interacts with, and its centrosomal localization is restricted by, CEP63. Protein levels of DTX4 were unaltered in *CEP63* depleted or ZIKV infected cells (Supplemental Figure 5N-O). Thus, we hypothesized that inhibition of CEP63 function increases centrosomal DTX4 which, in turn, targets TBK1 for K48 ubiquitin-mediate degradation.

To test this hypothesis, we immunoprecipitated TBK1 from cells lacking Cep63 or infected with ZIKV and immunoblotted for K48-linked ubiquitin chains (**Figure 5H-I** and Supplemental Figure 5P-Q). Like K63-ubiquitin, there was a marked increase in K48-ubiquitylated TBK1 in *Cep63* mutant and ZIKV-infected cells suggesting that increased centrosomal MIB1 and DTX4 impacts the K63 and K48-ubiquitylation and degradation of TBK1. These findings suggest that ZIKV disrupts centrosomal CEP63 altering the ubiquitylation and stability of TBK1 to dampen the innate immune response.

## Discussion

We have found that ZIKV-produced NS3 binds CEP63 to alter centrosome organization, but not centrosome amplification ^47^, increases centrosomal MIB1 and DTX4, decreases TBK1 levels, and disrupts the innate immune response. As CEP63 mutations cause microcephaly in humans ^7^, disrupted CEP63 function may underlie the pathogenesis of ZIKV-associated microcephaly. Moreover, the centrosomal CEP63 interactors MIB1 and DTX4, regulators of neurogenic NOTCH and innate immune signaling, respectively are altered upon either loss of CEP63 or ZIKV infection. Based on these results, we propose that ZIKV NS3 disrupts CEP63 function to both dampen the innate immune response and to disrupt developmental signaling during brain development.

Previous studies have reported conflicting effects on centrosome biogenesis in ZIKV infected cells ^16,17,19,20,47^. Our results using human neural progenitor cells from fetal tissue, induced pluripotent stem cells, and established neural cell lines indicate that acute ZIKV infection disrupts centrosome organization but does not lead to centrosome amplification. As studies by others on centrosome biogenesis examined later time points following ZIKV infection, centrosomal overabundance in ZIKV-infected cells may be secondary to multiple rounds of abnormal cell division.

We found that ZIKV-produced NS3 localizes to the centrosome to induce the formation of supernumerary Centrin foci by interacting with and localizing increased amounts of CEP63 to the centrosome. The failure of ZIKV infection in *Cep63* mutant MEFs to induce supernumerary Centrin, supports our assertion that ZIKV disrupts centrosome organization in a CEP63-dependent manner.

We and others have shown that CEP63 promotes centriole duplication ^7,8,11,48^. However, its function in other cellular processes has not been explored. Here, we have provided evidence that CEP63 controls TBK1 stability, a central component of innate immune signaling. Given that CEP63 interacts with and limits the localization of the centrosomal ubiquitin ligases, MIB1 and DTX4, to the centrosome, a direct role for CEP63 in promoting cellular signaling at the centrosome is implicit.

MIB1, a K63-linked ubiquitin ligase is a key regulator of NOTCH signaling, a pathway that controls neural progenitor maintenance and cell division ^49^. A less studied role for MIB1, is in the innate immune response ^21^. We have found that MIB1 over accumulates at the centrosome and K63-ubiquitylates TBK1 in response to ZIKV infection or CEP63 loss. The increased activity of MIB1 in ZIKV-infected neural progenitors or *CEP63*-depleted cells suggests that MIB1 activity in the NOTCH signaling pathway could also be affected. In agreement with this, a recent publication has demonstrated that MIB1 levels are affected in response to ZIKV infection ^18,50^, raising the possibility that ZIKV-associated microcephaly is a side effect of ZIKV altering the centrosome to evade host immunity.

We found that DTX4 localizes to the centrosome and promotes the K48-ubiquitylation of TBK1. Similar to MIB1, DTX4 accumulates at the centrosome in ZIKV infected cells in a CEP63-dependent manner. As TBK1 is ubiquitylated by DTX4 to promote its degradation, we propose that ZIKV mediated recruitment of DTX4 to the centrosome may limit TBK1 activity and stability, and thus the innate immune response, to ZIKV infection. It will be interesting to determine whether other viruses with NS3 homologues such as SARS-CoV-2 M^Pro^ and NSP13 which interact with centrosome proteins ^51^ can similarly suppress innate immunity by altering centrosome organization.

In summary, we have found that ZIKV NS3 localizes to the centrosome, recruits CEP63 and its binding partners MIB1 and DTX4 to ubiquitylate and degrade TBK1, a key regulator of the innate immune response. These findings provide mechanistic insight into how ZIKV specifically targets the centrosome with implications for how it may both evade viral detection and alter brain development.

## Materials and methods

### Cell culture and transfection

HeLa and 293T/17 cells (UCSF tissue culture facility) were cultured in Advanced Dulbecco’s Modified Eagle’s medium (DMEM, Thermo Fisher Scientific) supplemented with 2% fetal bovine serum (FBS, Thermo Fisher Scientific) and Glutamax-I (Thermo Fisher Scientific). U87 and H4 cells were cultured in DMEM (Thermo Fisher Scientific) supplemented with 10% FBS and L-glutamine (Thermo Fisher Scientific). *Cep63^gt/gt^* MEFs and control *Cep90^+/-^* MEFs were cultured in Amniomax C-100 (Thermo Fisher Scientific). 293T/17 and HeLa cells were transfected with plasmids using FugeneHD (Promega) or Lipofectamine3000 (Thermo Fisher Scientific), respectively, according to manufacturer’s instructions and analyzed 8 h later. NPCs were derived from pluripotent stem cells (NIH Human Embryonic Stem Cell Registry line WA09 (H9) at passages 30-35) according to a recently published protocol ^52^ and maintained in neural media composed of DMEM/F12 with sodium pyruvate and Glutamax, N2, B27, heparin and antibiotics. Medium was either supplemented with growth factors epidermal growth factor (10 ng/ml) and fibroblast growth factor (10 ng/ml).

### Fetal tissue collection, dissociation and culture

De-identified fetal brain tissue samples were collected with previous patient consent in strict observance of the legal and institutional ethical regulations from elective pregnancy termination specimens at San Francisco General Hospital. Protocols were approved by the Human Gamete, Embryo and Stem Cell Research Committee (an institutional review board) at the University of California, San Francisco. Blocks of cortical tissue spanning the ventricle to the cortical plate were dissected away from meninges and germinal zone using a stereomicroscope, and then minced using a razor blade. Cells were dissociated by incubation with Papain (Worthington Biochemical Corporation) at 37°C for 30-40 min, followed by addition of DNAse I and trituration. The cells were collected by centrifugation for 5 min at 300g, the supernatant was removed, and the cells were resuspended in sterile DMEM containing N-2, B-27 supplement, penicillin, streptomycin and glutamine and sodium pyruvate (0.11mg/mL) (all Invitrogen). The suspension was passed through a 40 μm strainer (BD Falcon) to yield a uniform suspension of single cells. Cells were plated at a density of 1.5×106 cells/well on 18mm coverslips (Neuvitro 18-GG-PDL) precoated with high concentration growth factor-reduced Matrigel (BDBiosciences, 354263) and cultured at 37°C, 5% CO_2_, 8% O_2_.

### *Cep90^-/-^* MEFs

We acquired *Cep90*^tm1.1(KOMP)VlCg^ (also called *Pibf1*^tm1.1(KOMP)VlCg^) heterozygous mice (Jackson Laboratory). *Cep90*^tm1.1(KOMP)VlCg^ is a deletion of Chr14:99099433-99254493 (GRCm38.p3), covering all 17 coding exons of *Cep90*. We isolated MEFs from littermate E8.5 embryos produced from a heterozygous intercross. MEFs were genotyped by quantitative PCR using EXPRESS SYBR GreenER qPCR Supermix, with premixed ROX (Invitrogen) on a 7300 Real-Time PCR machine (Applied Biosystems) and *Cep90* genotyping primers listed in the reagents table.

### Virus and infections

ZIKV strain PRVABC59 was propagated in Vero cells. Viral titers were determined by focus assay ^53^. Briefly, serial dilutions of viral stock were added to Vero cells in 96-well plates. 24 h post-infection, inoculum was removed and cells were fixed with 3.7% PFA for 15 minutes. Foci were visualized by immunofluorescent staining for flavivirus envelope. ZIKV infections of NPCs, U87 and H4 cell lines, at MOIs of 10, were carried out by incubating cells with inoculum for 1 h and then replacing the inoculum with fresh media. ZIKV was added to dissociated fetal brain cells at MOIs of 10 and incubated for 16 h unless stated otherwise. Mock and ZIKV-infected cells were fixed 12-16 h post infection in chilled methanol for 3 min at −20°C and processed for immunofluorescence.

### Molecular Biology

Wild type human codon optimized ZIKV NS2B3 and NS3 open reading frame flanked by attB sites were synthesized by Gibson assembly (SGI). Gateway cloning into pDONR221 generated pENTR-NS3 ZIKV. Subsequent Gateway-mediated subcloning into pDEST-CMV-Myc (gift of Keith Yamamoto) generated pCMV-Myc-NS3 Brazilian and Ugandan ZIKV, encoding N-terminally Myc-tagged NS3 and pcMV-Myc-NS2B3 Brazilian ZIKV encoding an N-terminally Myc-tagged fusion of NS2B and NS3. The S135A mutant form of NS3 was generated using site-directed mutagenesis to create pCMV-Myc-NS3-S135A Brazilian ZIKV (QuikChange II, Agilent).

### Biochemistry

Immunoprecipitations were performed as previously described ^54^. In brief, mock or ZIKV infected U87 cells or 293T/17 cells were collected in Dulbecco’s phosphate-buffered saline (DPBS), lysed in lysis buffer (1% IGEPAL CA-630, 50mM Tris-HCL pH7.4, 266mM NaCl, 2.27mM KCl, 1.25mM KH_2_PO_4_, 6.8mM Na_2_HPO_4_-7H_2_O) supplemented with protease and phosphatase inhibitors (EMD Millipore). Myc-tagged proteins were immunoprecipitated with 4AG Myc monoclonal agarose beads (EMD Millipore), washed three times in lysis buffer and boiled in 2X Laemmli reducing buffer (Bio-Rad). Samples were separated on 4-15% TGX precast gels (Bio-Rad), transferred onto Protran BA85 nitrocellulose membrane (GE Healthcare) and subsequently analyzed by immunoblot using ECL Lightening PLUS (Perkin-Elmers) or SuperSignal West Dura (Thermo Fisher Scientific).

### Immunofluorescence microscopy

Cells were fixed in −20°C methanol for 3 minutes followed by permeabilization in blocking buffer (2.5% BSA, 0.1% Triton-X100, 0.03% NaN_3_ in DPBS) for 30 min. Primary and secondary antibodies were diluted in blocking buffer and incubated with cells for at least 1 h. To detect cells in S-phase, cells were co-stained with antibodies to Centrin and Cyclin A to determine centriole number and S-phase/G2 cells, respectively. To detect Phospho-TBK1, U87 and MEFs were fixed in 4% paraformaldehyde for 30 minutes, blocked in 5% FBS and 0.5% TritonX100 diluted in DPBS. Permeabilized cells were stained overnight at room temperature using the p-TBK1 antibody at a dilution of 1:50 in 5% FBS and 0.1% TritonX100 diluted in DPBS. Samples were mounted in Gelvatol and imaged with an Axio Observer D1 or LSM700 (Zeiss). Images were processed using Adobe Photoshop and analyzed using Fiji.

### RT-ddPCR

RNA was isolated from infected cells at various time points post infection using Trizol (Invitrogen) and Direct-zol RNA Miniprep kits (Zymo Research). Droplets containing RNA, One-Step RT-ddPCR Advanced Kit for Probes (Bio-Rad Laboratories), and probes were prepared on a QX200 Droplet Generator (Bio-Rad Laboratories), using Droplet Generation Oil for Probes (Bio-Rad Laboratories). RT-PCR was run in droplets on a C1000 Touch (Bio-Rad Laboratories) following the manufactures direction for One-Step RT-ddPCR Advanced Kit for Probes. Droplets were read on a QX100 Droplet Reader (Bio-Rad Laboratories). Primers: Forward IFNβ 5’-GATGACGGAGAAGATGCAGAAG-3’, Reverse IFNβ 5’-ACCCAGTGCTGGAGAAATTG-3’. The probe was designed using the Primer Quest Tool from IDT and used 5’6’FAM/ZEN/3’IBFQ (IDT): 5’-/5HEX/ACACTGCCT/ZEN/TTGCCATCCAAGAGA/3IABkFQ/-3’.

#### Antibodies used

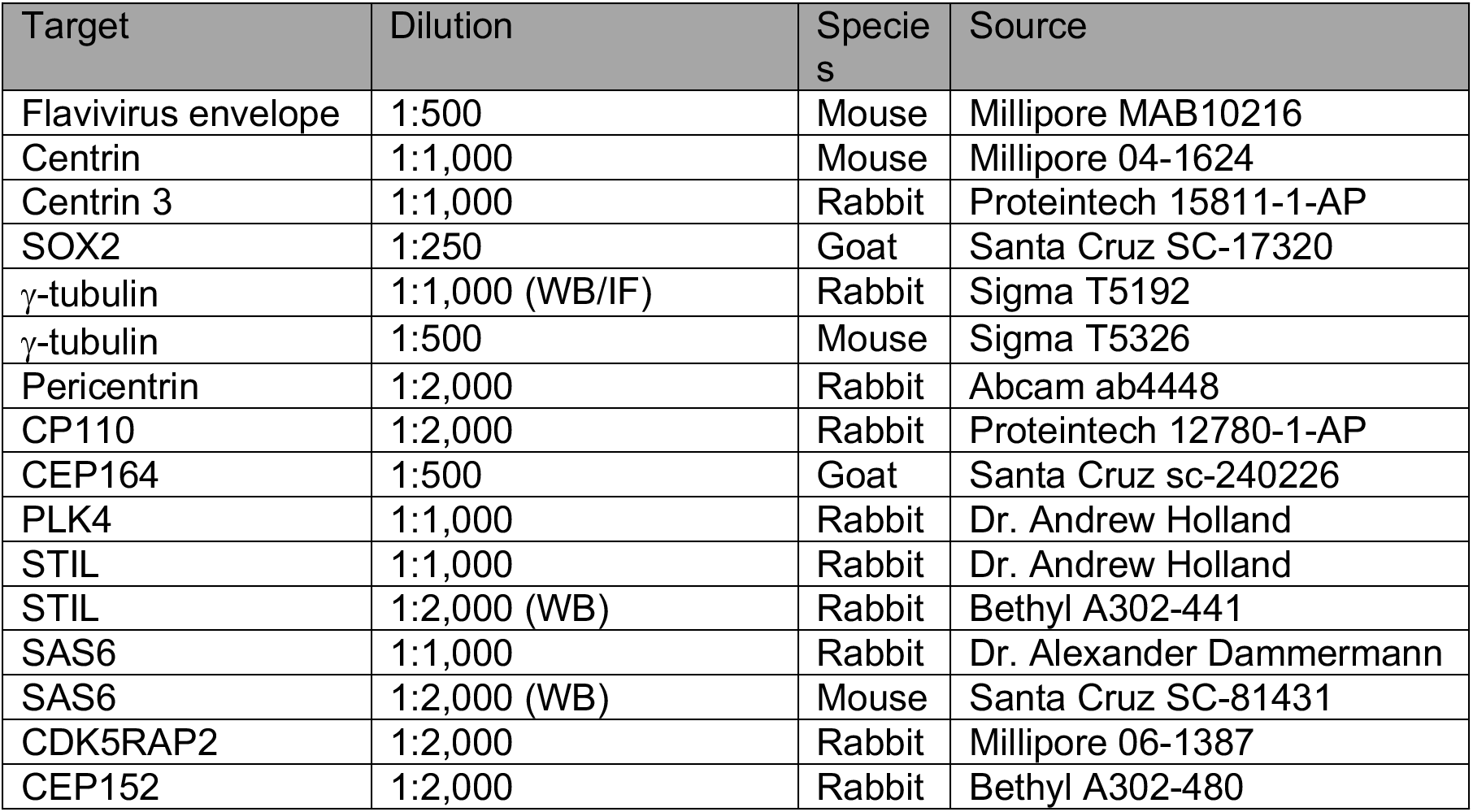
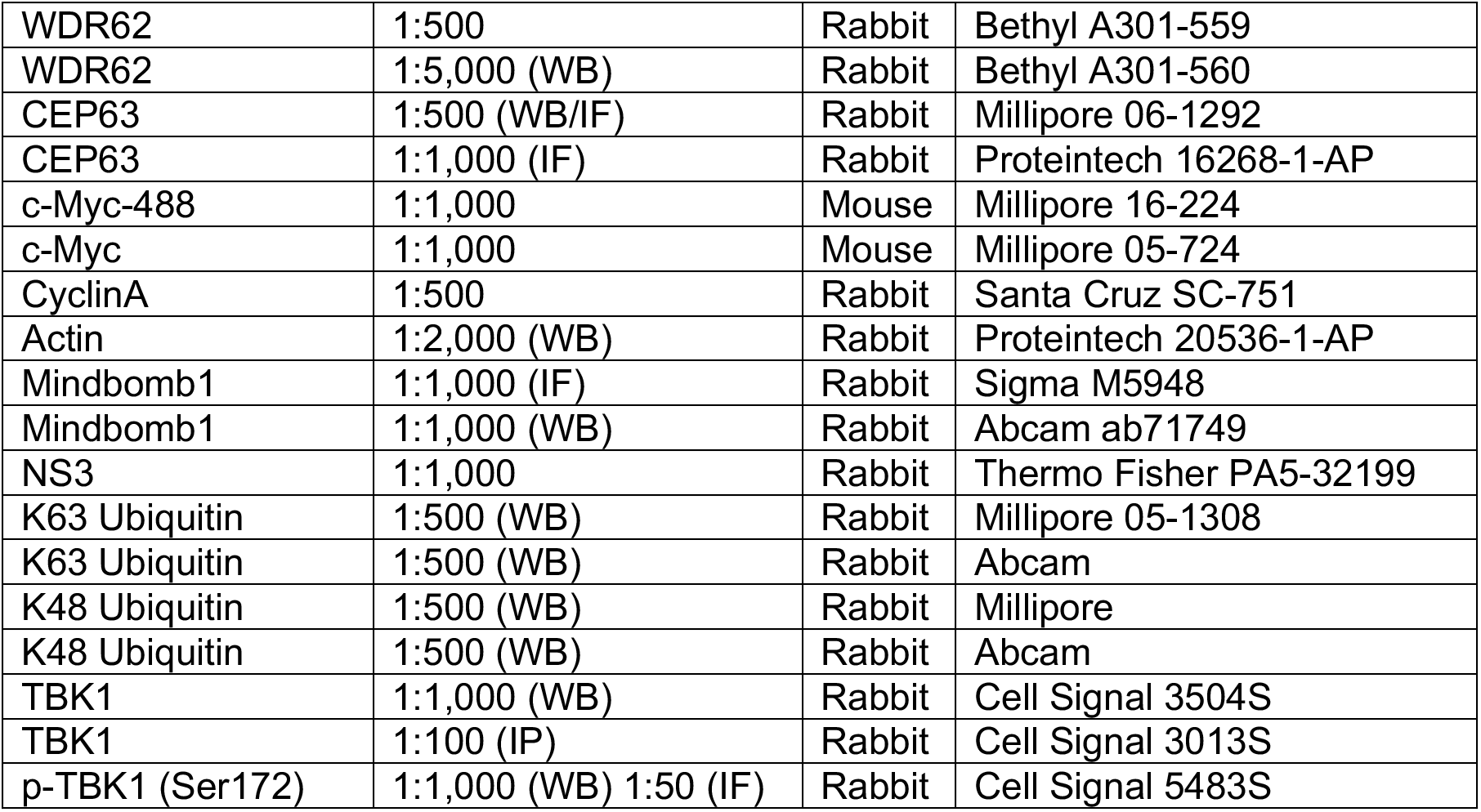

Antibodies used:
| Target | Dilution | Species | Source |
| --- | --- | --- | --- |
| Flavivirus envelope | 1:500 | Mouse | Millipore MAB10216 |
| Centrin | 1:1,000 | Mouse | Millipore 04-1624 |
| Centrin 3 | 1:1,000 | Rabbit | Proteintech 15811-1-AP |
| SOX2 | 1:250 | Goat | Santa Cruz SC-17320 |
| γ-tubulin | 1:1,000 (WB/IF) | Rabbit | Sigma T5192 |
| γ-tubulin | 1:500 | Mouse | Sigma T5326 |
| Pericentrin | 1:2,000 | Rabbit | Abcam ab4448 |
| CP110 | 1:2,000 | Rabbit | Proteintech 12780-1-AP |
| CEP164 | 1:500 | Goat | Santa Cruz sc-240226 |
| PLK4 | 1:1,000 | Rabbit | Dr. Andrew Holland |
| STIL | 1:1,000 | Rabbit | Dr. Andrew Holland |
| STIL | 1:2,000 (WB) | Rabbit | Bethyl A302-441 |
| SAS6 | 1:1,000 | Rabbit | Dr. Alexander Dammermann |
| SAS6 | 1:2,000 (WB) | Mouse | Santa Cruz SC-81431 |
| CDK5RAP2 | 1:2,000 | Rabbit | Millipore 06-1387 |
| CEP152 | 1:2,000 | Rabbit | Bethyl A302-480 |
| WDR62 | 1:500 | Rabbit | Bethyl A301-559 |
| WDR62 | 1:5,000 (WB) | Rabbit | Bethyl A301-560 |
| CEP63 | 1:500 (WB/IF) | Rabbit | Millipore 06-1292 |
| CEP63 | 1:1,000 (IF) | Rabbit | Proteintech 16268-1-AP |
| c-Myc-488 | 1:1,000 | Mouse | Millipore 16-224 |
| c-Myc | 1:1,000 | Mouse | Millipore 05-724 |
| CyclinA | 1:500 | Rabbit | Santa Cruz SC-751 |
| Actin | 1:2,000 (WB) | Rabbit | Proteintech 20536-1-AP |
| Mindbomb1 | 1:1,000 (IF) | Rabbit | Sigma M5948 |
| Mindbomb1 | 1:1,000 (WB) | Rabbit | Abcam ab71749 |
| NS3 | 1:1,000 | Rabbit | Thermo Fisher PA5-32199 |
| K63 Ubiquitin | 1:500 (WB) | Rabbit | Millipore 05-1308 |
| K63 Ubiquitin | 1:500 (WB) | Rabbit | Abcam |
| K48 Ubiquitin | 1:500 (WB) | Rabbit | Millipore |
| K48 Ubiquitin | 1:500 (WB) | Rabbit | Abcam |
| TBK1 | 1:1,000 (WB) | Rabbit | Cell Signal 3504S |
| TBK1 | 1:100 (IP) | Rabbit | Cell Signal 3013S |
| p-TBK1 (Ser172) | 1:1,000 (WB) 1:50 (IF) | Rabbit | Cell Signal 5483S |

#### Primers used

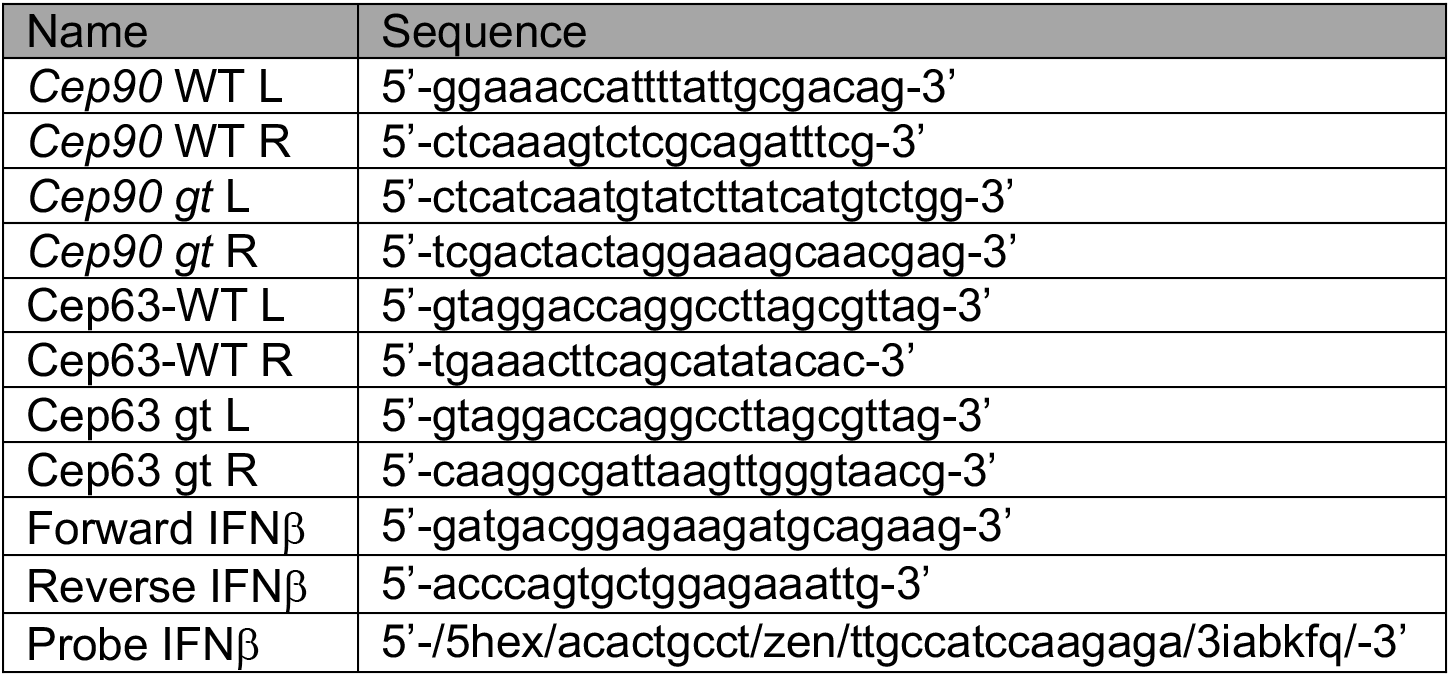

---

**Supplemental Figure 1:**
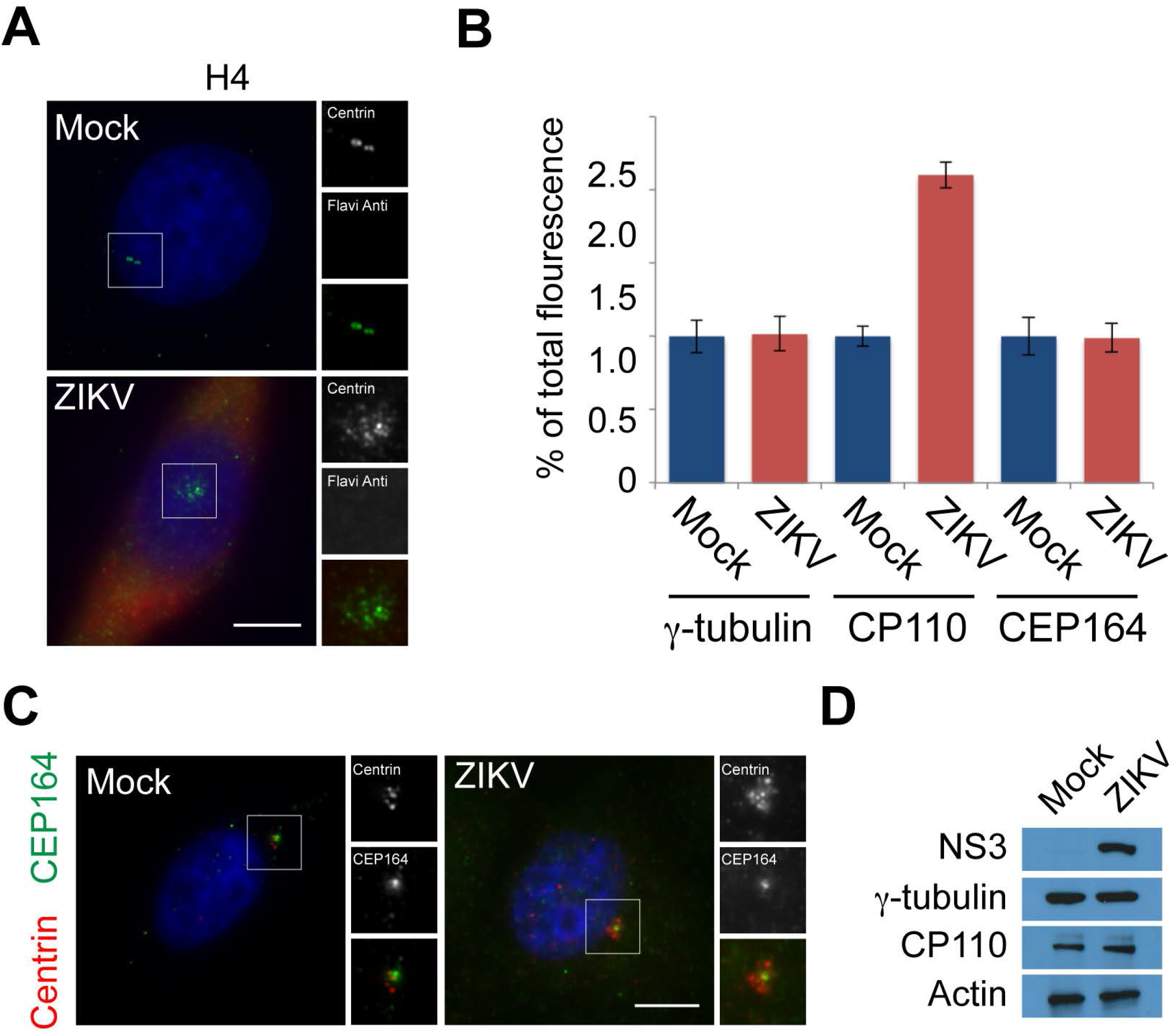
**(A)** Mock and ZIKV-infected H4 cells co-stained for Centrin (green) and flavivirus envelope (red) to mark centrioles and infected cells, respectively. All cells were fixed 16hpi. Scale bars indicate 5μm for all images. **(B)** Quantification of mean fluorescence intensities ± s.d. of γ-tubulin, CP110, and CEP164 in mock and ZIKV infected U87 cells. For fluorescence quantifications 8 cells were analyzed per experiment (n=3). Asterisk denotes p<0.005 (Student’s t test). **(C)** Immunofluorescence analysis of mock and ZIKV-infected U87 cells co-stained for Centrin (red) and CEP164 (green). Scale bars indicate 5μM for all images. **(D)** Immunoblot of mock or ZIKV-infected U87 cell lysates probed for NS3, CP110, or γ-tubulin. Actin served as a loading control.

**Supplemental Figure 2:**
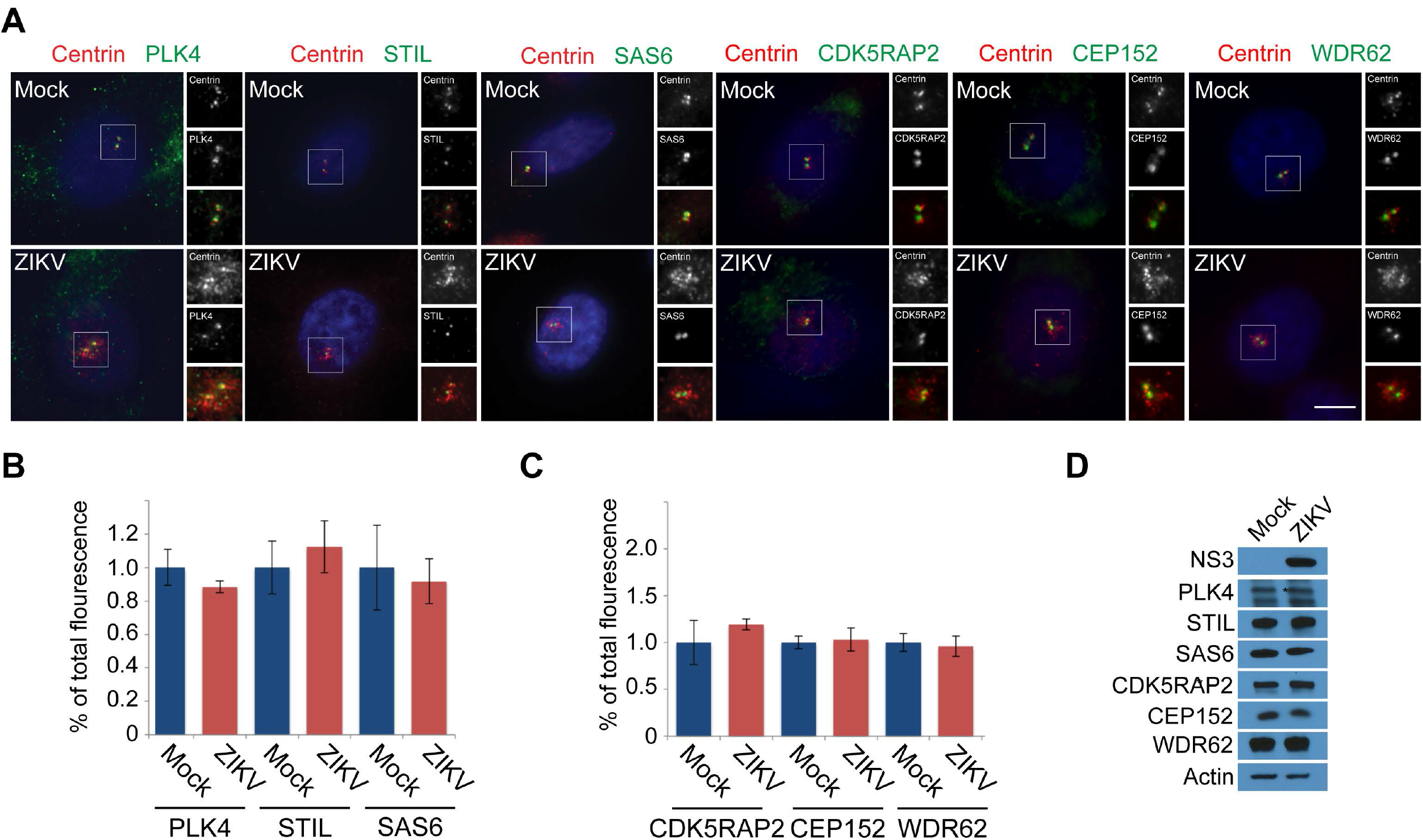
**(A)** Immunofluorescence analysis of S phase mock and ZIKV-infected U87 cells costained with antibodies to Centrin (red), and the microcephaly-associated centriole duplication factors: PLK4 (green), STIL (green), SAS6 (green), CDK5RAP2 (green), CEP152 (green) or WDR62 (green). **(B-C)** The fluorescence intensities ± s.d. of PLK4, STIL, SAS6, CDK5RAP2, CEP152 and WDR62 were quantified in mock and ZIKV infected U87 cells. For fluorescence quantifications 8 cells were analyzed per experiment (n=3). Asterisk denotes p<0.005 (Student’s t test). **(D)** Immunoblot of mock or ZIKV-infected U87 cell lysates probed for NS3, PLK4, STIL, SAS6, CDK5RAP2, CEP152, and WDR62. Actin served as a loading control.

**Supplemental Figure 3:**
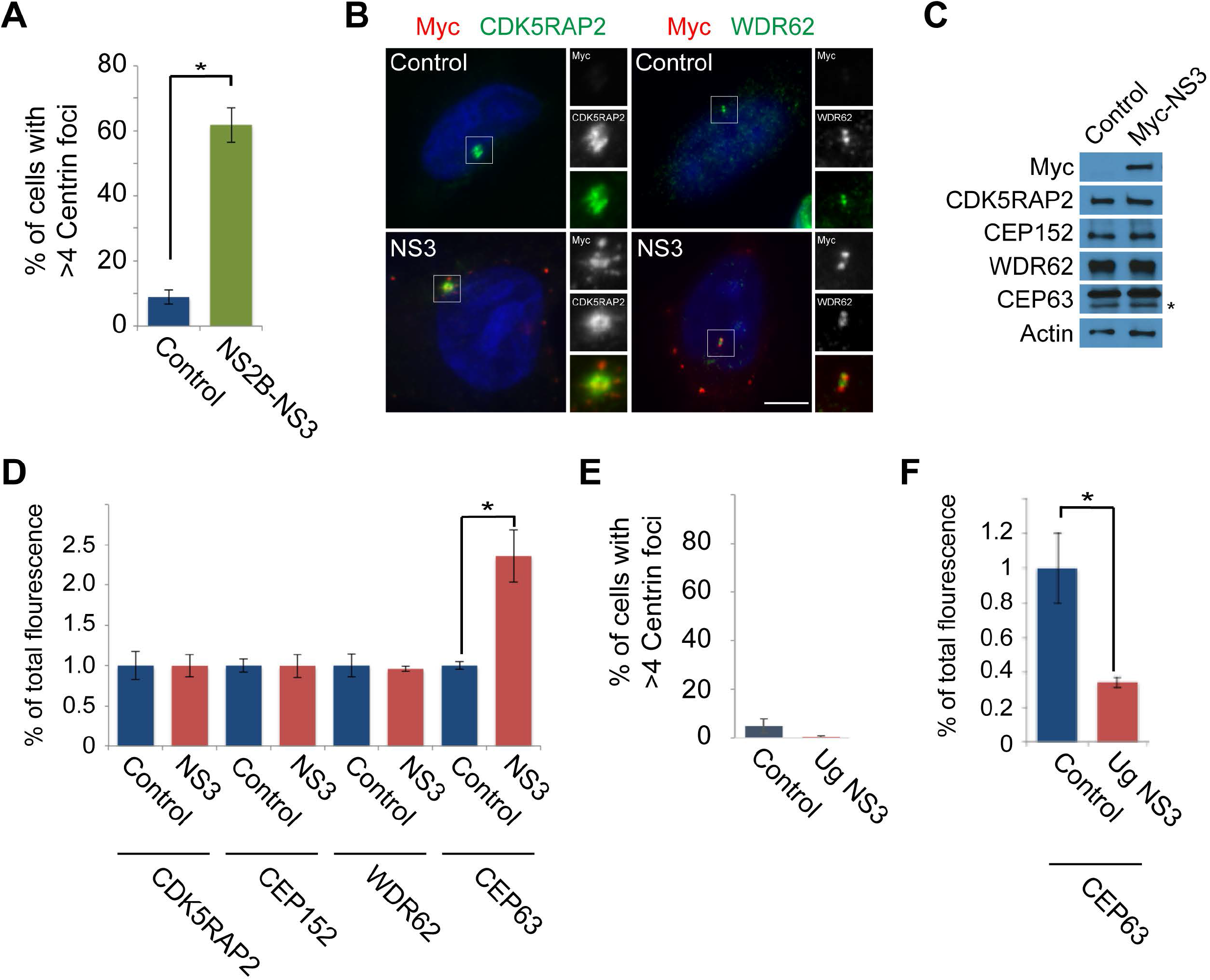
**(A)** Quantification of the percentage of control or NS2B-NS3 expressing cells in S-phase with greater than four Centrin foci. Asterisk denotes p<0.005 (Student’s t test). **(B)** Control or Myc-tagged Brazilian ZIKV NS3 (Br NS3) were co-stained for Myc (red) and CDK5RAP2 (green) or WDR62 (green). Scale bars indicate 5μm for all images. **(C)** Immunoblot of HeLa cells expressing Myc-tagged NS3 probed for c-Myc, CDK5RAP2, CEP152, WDR62, and CEP63. Actin served as a loading control. Asterisk denotes specific band for CEP63. **(D)** Quantification of mean fluorescence intensities ± s.d. of CDK5RAP2, CEP152, WDR62, and CEP63 in HeLa cells transfected with an empty c-Myc vector (Control) or Brazilian Myc-NS3. For fluorescence quantifications 8 cells were analyzed per experiment (n=3). Asterisk denotes p<0.005 (Student’s t test). **(E)** Quantification of HeLa cells transfected with Ugandan Myc-NS3 with greater than four centrioles. **(F)** Quantification of mean fluorescence intensities ± s.d. of CEP63 in control and Ugandan Myc-NS3 expressing HeLa cells.

**Supplemental Figure 4:**
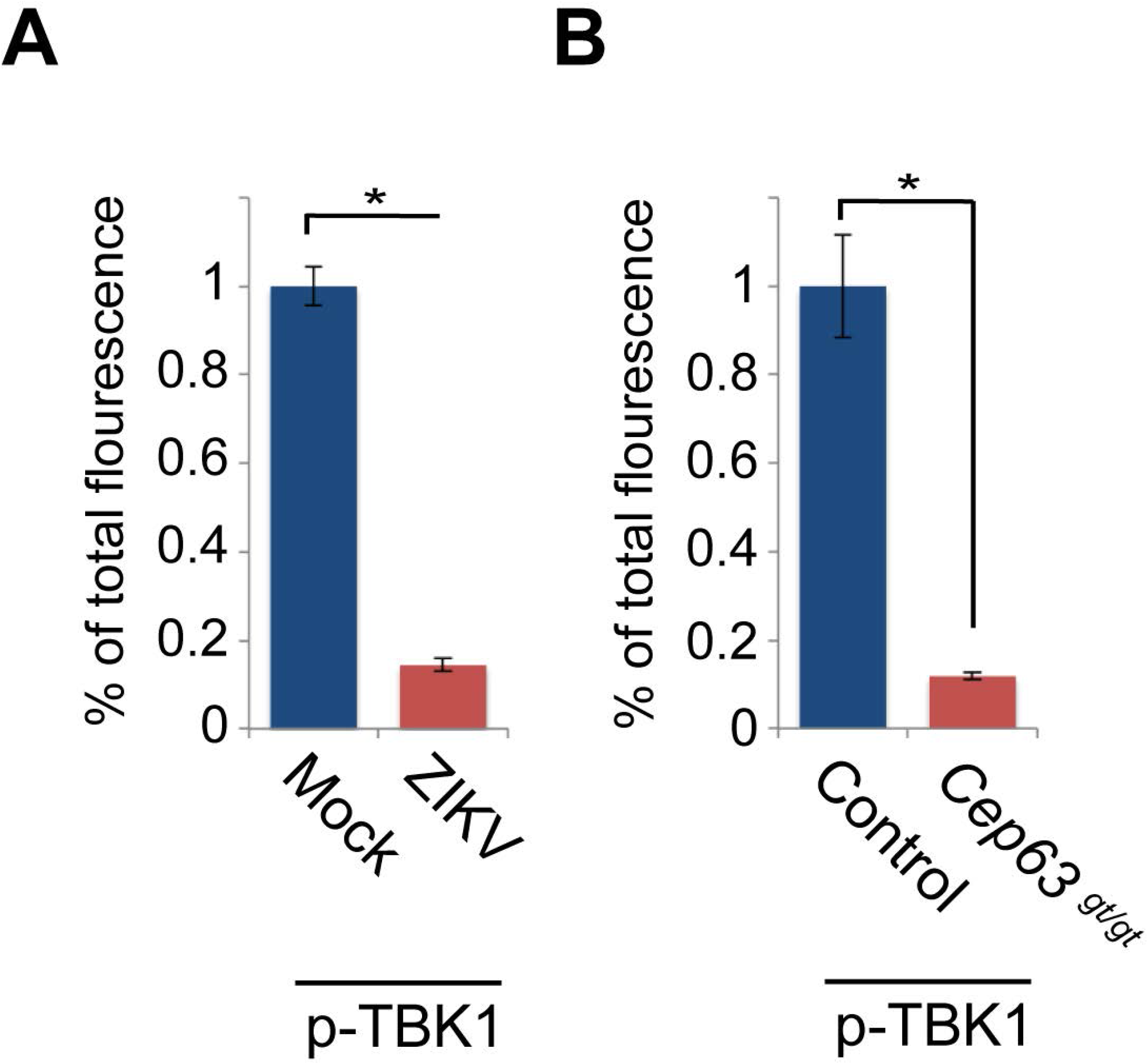
**(A)** Quantification of mean fluorescence intensities ± s.d. of centrosomal p-TBK1 in mock and ZIKV-infected U87 cells. For fluorescence quantifications 8 cells were analyzed per experiment (n=3). Asterisk denotes p<0.005 (Student’s t test). **(B)** Mean fluorescence intensity quantifications of centrosomal phospho-Tbk1 (p-Tbk1) in Control and *Cep63*^gt/gt^ MEFs.

**Supplemental Figure 5:**
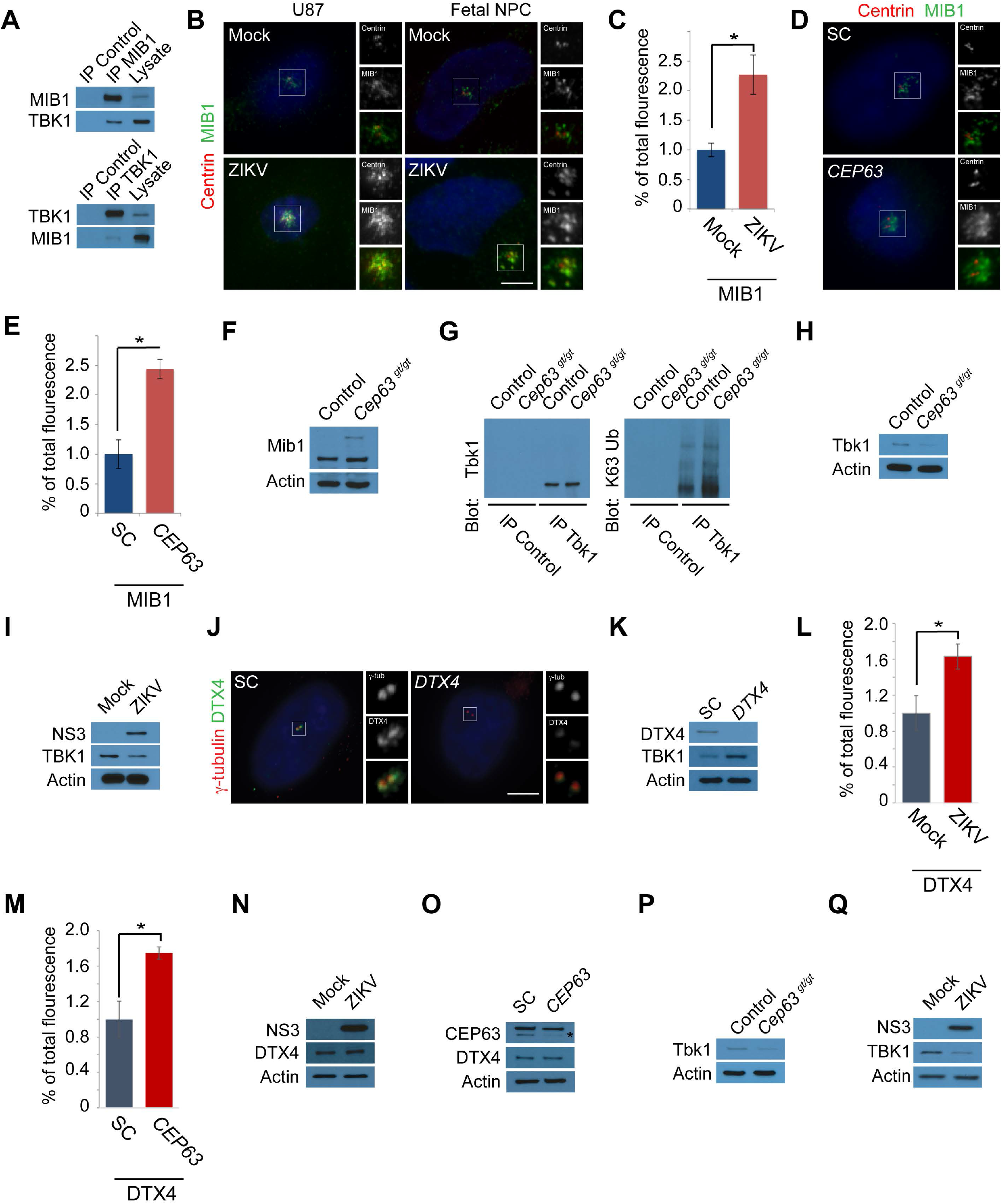
**(A)** HeLa cell lysate was subjected to immunoprecipitation using antibodies to MIB1, TBK1 and c-Myc (negative control). Coprecipitated proteins were analyzed by western blotting using antibodies to MIB1 and TBK1. **(B)** Mock or ZIKV infected fetal neural precursors and U87 cells co-stained for Centrin (red) and MIB1 (green). All cells were fixed and stained 16 hpi. Scale bars indicate 5μM for all images. **(C)** Quantification of mean fluorescence intensities ± s.d. of MIB1 in mock and ZIKV-infected U87 cells. For fluorescence quantifications 8 cells were analyzed per experiment (n=3). Asterisk denotes p<0.005 (Student’s t test). **(D)** S phase SC and *CEP63* siRNA transfected HeLa cells co-stained for Centrin (red) and MIB1 (green). **(E)** Mean fluorescence intensity quantifications of MIB1 in SC and *CEP63* siRNA-treated HeLa cells. **(F)** Lysates from Control and *Cep63*^gt/gt^ MEFs immunoblotted for Mib1. Asterisk denotes a higher molecular weight species of Mib1. Actin served as a loading control. **(G)** Tbk1 was immunoprecipitated from Control and *Cep63^gt/gt^* MEF total cell lysate. Precipitating proteins were probed for Tbk1 or K63-linked ubiquitin. **(H)** Cell lysates from Control and *Cep63^gt/gt^* MEFs used in immunoprecipitation experiments in Supplemental Figure 5G. Lysates were probed with an antibody to Tbk1. Actin served as a loading control. **(I)** Mock and ZIKV-infected U87 cell lysates used to immunoprecipitate TBK1 in Figure 5D were analyzed by western blot using antibodies to NS3 and TBK1. Actin served as a loading control. **(J)** SC or *DTX4*-depleted HeLa cells were co-stained for γ-tubulin and DTX4. **(K)** Immunoblot of HeLa cells transfected with SC or *DTX4* siRNA probed for DTX4 and TBK1. Actin served as a loading control. **(L)** Quantification of mean fluorescence intensities ± s.d. of DTX4 in mock or ZIKV-infected U87 cells. **(M)** Quantification of centrosomal DTX4 in SC and *CEP63* depleted HeLa cells expressed as a mean fluorescence intensities ± s.d. of the control. **(N)** Total cell lysate from mock and ZIKV-transfected U87 cells 16 hpi were analyzed by western blot using antibodies to ZIKV NS3 and DTX4. Actin served as a loading control. **(O)** Immunoblot analysis of lysate from SC and *CEP63* siRNA-treated HeLa cells using antibodies to CEP63 and DTX4. Actin served as a loading control. **(P)** Control and *Cep63^gt/gt^* MEF lysates used to immunoprecipitate Tbk1 in Figure 5M were analyzed by western blot using antibodies to Tbk1. Actin served as a loading control. **(Q)** Cell lysates from mock and ZIKV-infected U87 cells used to immunoprecipitate TBK1 in Figure 5N. Lysates were probed with an antibody to ZIKV NS3 and TBK1. Actin served as a loading control.

## Bibliography

1 Jayaraman, D., Bae, B. I. & Walsh, C. A. The Genetics of Primary Microcephaly. Annual review of genomics and human genetics 19, 177–200, doi:10.1146/annurev-genom-083117-021441 (2018).

2 Saade, M., Blanco-Ameijeiras, J., Gonzalez-Gobartt, E. & Marti, E. A centrosomal view of CNS growth. Development 145, doi:10.1242/dev.170613 (2018).

3 Nano, M. & Basto, R. Consequences of Centrosome Dysfunction During Brain Development. Adv Exp Med Biol 1002, 19–45, doi:10.1007/978-3-319-57127-0_2 (2017).

4 Bertipaglia, C., Goncalves, J. C. & Vallee, R. B. Nuclear migration in mammalian brain development. Semin Cell Dev Biol 82, 57–66, doi:10.1016/j.semcdb.2017.11.033 (2018).

5 Gonczy, P. & Hatzopoulos, G. N. Centriole assembly at a glance. J Cell Sci 132, doi:10.1242/jcs.228833 (2019).

6 Varadarajan, R. & Rusan, N. M. Bridging centrioles and PCM in proper space and time. Essays Biochem 62, 793–801, doi:10.1042/EBC20180036 (2018).

7 Sir, J. H. et al. A primary microcephaly protein complex forms a ring around parental centrioles. Nat Genet 43, 1147–1153, doi:10.1038/ng.971 (2011).

8 Brown, N. J., Marjanovic, M., Luders, J., Stracker, T. H. & Costanzo, V. Cep63 and cep152 cooperate to ensure centriole duplication. PLoS One 8, e69986, doi:10.1371/journal.pone.0069986 (2013).

9 Kim, T. S. et al. Molecular architecture of a cylindrical self-assembly at human centrosomes. Nature communications 10, 1151, doi:10.1038/s41467-019-08838-2 (2019).

10 Jayaraman, D. et al. Microcephaly Proteins Wdr62 and Aspm Define a Mother Centriole Complex Regulating Centriole Biogenesis, Apical Complex, and Cell Fate. Neuron 92, 813–828, doi:10.1016/j.neuron.2016.09.056 (2016).

11 Kodani, A. et al. Centriolar satellites assemble centrosomal microcephaly proteins to recruit CDK2 and promote centriole duplication. eLife 4, doi:10.7554/eLife.07519 (2015).

12 Sonnen, K. F., Gabryjonczyk, A. M., Anselm, E., Stierhof, Y. D. & Nigg, E. A. Human Cep192 and Cep152 cooperate in Plk4 recruitment and centriole duplication. J Cell Sci 126, 3223–3233, doi:10.1242/jcs.129502 (2013).

13 Lawo, S., Hasegan, M., Gupta, G. D. & Pelletier, L. Subdiffraction imaging of centrosomes reveals higher-order organizational features of pericentriolar material. Nat Cell Biol 14, 1148–1158, doi:10.1038/ncb2591 (2012).

14 Hu, W. F. et al. Katanin p80 regulates human cortical development by limiting centriole and cilia number. Neuron 84, 1240–1257, doi:10.1016/j.neuron.2014.12.017 (2014).

15 Bazzi, H. & Anderson, K. V. Acentriolar mitosis activates a p53-dependent apoptosis pathway in the mouse embryo. Proc Natl Acad Sci U S A 111, E1491–1500, doi:10.1073/pnas.1400568111 (2014).

16 Wolf, B. et al. Zika virus causes supernumerary foci with centriolar proteins and impaired spindle positioning. Open Biol 7, doi:10.1098/rsob.160231 (2017).

17 Souza, B. S. et al. Zika virus infection induces mitosis abnormalities and apoptotic cell death of human neural progenitor cells. Sci Rep 6, 39775, doi:10.1038/srep39775 (2016).

18 Ferraris, P. et al. Zika virus differentially infects human neural progenitor cells according to their state of differentiation and dysregulates neurogenesis through the Notch pathway. Emerging microbes & infections 8, 1003–1016, doi:10.1080/22221751.2019.1637283 (2019).

19 Gabriel, E. et al. Recent Zika Virus Isolates Induce Premature Differentiation of Neural Progenitors in Human Brain Organoids. Cell stem cell 20, 397–406 e395, doi:10.1016/j.stem.2016.12.005 (2017).

20 Kesari, A. S. et al. Zika virus NS5 localizes at centrosomes during cell division. Virology 541, 52–62, doi:10.1016/j.virol.2019.11.018 (2020).

21 Li, S., Wang, L., Berman, M., Kong, Y. Y. & Dorf, M. E. Mapping a dynamic innate immunity protein interaction network regulating type I interferon production. Immunity 35, 426–440, doi:10.1016/j.immuni.2011.06.014 (2011).

22 Fitzgerald, K. A. et al. IKKepsilon and TBK1 are essential components of the IRF3 signaling pathway. Nature immunology 4, 491–496, doi:10.1038/ni921 (2003).

23 Liu, D. et al. SOCS3 Drives Proteasomal Degradation of TBK1 and Negatively Regulates Antiviral Innate Immunity. Mol Cell Biol 35, 2400–2413, doi:10.1128/MCB.00090-15 (2015).

24 Perry, A. K., Chow, E. K., Goodnough, J. B., Yeh, W. C. & Cheng, G. Differential requirement for TANK-binding kinase-1 in type I interferon responses to toll-like receptor activation and viral infection. J Exp Med 199, 1651–1658, doi:10.1084/jem.20040528 (2004).

25 Wang, L., Li, S. & Dorf, M. E. NEMO binds ubiquitinated TANK-binding kinase 1 (TBK1) to regulate innate immune responses to RNA viruses. PLoS One 7, e43756, doi:10.1371/journal.pone.0043756 (2012).

26 Paoletti, A., Moudjou, M., Paintrand, M., Salisbury, J. L. & Bornens, M. Most of centrin in animal cells is not centrosome-associated and centrosomal centrin is confined to the distal lumen of centrioles. J Cell Sci 109 (Pt 13), 3089–3102 (1996).

27 Spektor, A., Tsang, W. Y., Khoo, D. & Dynlacht, B. D. Cep97 and CP110 suppress a cilia assembly program. Cell 130, 678–690, doi:10.1016/j.cell.2007.06.027 (2007).

28 Graser, S. et al. Cep164, a novel centriole appendage protein required for primary cilium formation. J Cell Biol 179, 321–330, doi:10.1083/jcb.200707181 (2007).

29 Stearns, T., Evans, L. & Kirschner, M. Gamma-tubulin is a highly conserved component of the centrosome. Cell 65, 825–836, doi:10.1016/0092-8674(91)90390-k (1991).

30 Khan, M. A. et al. A missense mutation in the PISA domain of HsSAS-6 causes autosomal recessive primary microcephaly in a large consanguineous Pakistani family. Human molecular genetics 23, 5940–5949, doi:10.1093/hmg/ddu318 (2014).

31 Kumar, A., Girimaji, S. C., Duvvari, M. R. & Blanton, S. H. Mutations in STIL, encoding a pericentriolar and centrosomal protein, cause primary microcephaly. Am J Hum Genet 84, 286–290, doi:10.1016/j.ajhg.2009.01.017 S0002-9297(09)00024-X [pii] (2009).

32 Martin, C. A. et al. Mutations in PLK4, encoding a master regulator of centriole biogenesis, cause microcephaly, growth failure and retinopathy. Nat Genet 46, 1283–1292, doi:10.1038/ng.3122 (2014).

33 Le Breton, M. et al. Flavivirus NS3 and NS5 proteins interaction network: a high-throughput yeast two-hybrid screen. BMC microbiology 11, 234, doi:10.1186/1471-2180-11-234 (2011).

34 Bera, A. K., Kuhn, R. J. & Smith, J. L. Functional characterization of cis and trans activity of the Flavivirus NS2B-NS3 protease. J Biol Chem 282, 12883–12892, doi:10.1074/jbc.M611318200 (2007).

35 Calvet, G. et al. Detection and sequencing of Zika virus from amniotic fluid of fetuses with microcephaly in Brazil: a case study. Lancet Infect Dis 16, 653–660, doi:10.1016/S1473-3099(16)00095-5 (2016).

36 Mlakar, J. et al. Zika Virus Associated with Microcephaly. N Engl J Med 374, 951–958, doi:10.1056/NEJMoa1600651 (2016).

37 Zhu, Z. et al. Comparative genomic analysis of pre-epidemic and epidemic Zika virus strains for virological factors potentially associated with the rapidly expanding epidemic. Emerging microbes & infections 5, e22, doi:10.1038/emi.2016.48 (2016).

38 Grant, A. et al. Zika Virus Targets Human STAT2 to Inhibit Type I Interferon Signaling. Cell host & microbe 19, 882–890, doi:10.1016/j.chom.2016.05.009 (2016).

39 Kumar, A. et al. Zika virus inhibits type-I interferon production and downstream signaling. EMBO Rep 17, 1766–1775, doi:10.15252/embr.201642627 (2016).

40 Sharma, S. et al. Triggering the interferon antiviral response through an IKK-related pathway. Science 300, 1148–1151, doi:10.1126/science.1081315 (2003).

41 Cui, J. et al. NLRP4 negatively regulates type I interferon signaling by targeting the kinase TBK1 for degradation via the ubiquitin ligase DTX4. Nat Immunol 13, 387–395, doi:10.1038/ni.2239 (2012).

42 Farlik, M. et al. Contribution of a TANK-binding kinase 1-interferon (IFN) regulatory factor 7 pathway to IFN-gamma-induced gene expression. Mol Cell Biol 32, 1032–1043, doi:10.1128/MCB.06021-11 (2012).

43 Helgason, E., Phung, Q. T. & Dueber, E. C. Recent insights into the complexity of Tank-binding kinase 1 signaling networks: the emerging role of cellular localization in the activation and substrate specificity of TBK1. FEBS Lett 587, 1230–1237, doi:10.1016/j.febslet.2013.01.059 (2013).

44 Onorati, M. et al. Zika Virus Disrupts Phospho-TBK1 Localization and Mitosis in Human Neuroepithelial Stem Cells and Radial Glia. Cell reports, doi:10.1016/j.celrep.2016.08.038 (2016).

45 Pillai, S. et al. Tank binding kinase 1 is a centrosome-associated kinase necessary for microtubule dynamics and mitosis. Nat Commun 6, 10072, doi:10.1038/ncomms10072 (2015).

46 Gupta, G. D. et al. A Dynamic Protein Interaction Landscape of the Human Centrosome-Cilium Interface. Cell 163, 1484–1499, doi:10.1016/j.cell.2015.10.065 (2015).

47 Onorati, M. et al. Zika Virus Disrupts Phospho-TBK1 Localization and Mitosis in Human Neuroepithelial Stem Cells and Radial Glia. Cell reports 16, 2576–2592, doi:10.1016/j.celrep.2016.08.038 (2016).

48 Loffler, H. et al. Cep63 recruits Cdk1 to the centrosome: implications for regulation of mitotic entry, centrosome amplification, and genome maintenance. Cancer Res 71, 2129–2139, doi:10.1158/0008-5472.CAN-10-2684 (2011).

49 Koo, B. K. et al. Mind bomb 1 is essential for generating functional Notch ligands to activate Notch. Development 132, 3459–3470, doi:10.1242/dev.01922 (2005).

50 Wen, F. et al. Zika virus increases mind bomb 1 levels, causing degradation of pericentriolar material 1 (PCM1) and dispersion of PCM1-containing granules from the centrosome. J Biol Chem 294, 18742–18755, doi:10.1074/jbc.RA119.010973 (2019).

51 Gordon, D. E. et al. A SARS-CoV-2-Human Protein-Protein Interaction Map Reveals Drug Targets and Potential Drug-Repurposing. bioRxiv, doi:10.1101/2020.03.22.002386 (2020).

52 Krencik, R. et al. Dysregulation of astrocyte extracellular signaling in Costello syndrome. Science translational medicine 7, 286ra266, doi:10.1126/scitranslmed.aaa5645 (2015).

53 Battegay, M. et al. Quantification of lymphocytic choriomeningitis virus with an immunological focus assay in 24-or 96-well plates. Journal of virological methods 33, 191–198 (1991).

54 Kodani, A., Salome Sirerol-Piquer, M., Seol, A., Garcia-Verdugo, J. M. & Reiter, J. F. Kif3a interacts with Dynactin subunit p150 Glued to organize centriole subdistal appendages. The EMBO journal 32, 597–607, doi:10.1038/emboj.2013.3 (2013).

